# Proteolytic queues at ClpXP increase antibiotic tolerance

**DOI:** 10.1101/680504

**Authors:** Heather S. Deter, Alawiah H. Abualrahi, Prajakta Jadhav, Elise K. Schweer, Curtis T. Ogle, Nicholas C. Butzin

## Abstract

Antibiotic tolerance is a widespread phenomenon that renders antibiotic treatments less effective and facilitates antibiotic resistance. Here we explore the role of proteases in antibiotic tolerance, short-term population survival of antibiotics, using queueing theory (i.e. the study of waiting lines), computational models, and a synthetic biology approach. Proteases are key cellular components that degrade proteins and play an important role in a multi-drug tolerant subpopulation of cells, called persisters. We found that queueing at the protease ClpXP increases antibiotic tolerance ~80 and ~60 fold in an *E. coli* population treated with ampicillin and ciprofloxacin, respectively. There does not appear to be an effect on antibiotic persistence, which we distinguish from tolerance based on population decay. These results demonstrate that proteolytic queueing is a practical method to probe bacterial tolerance and related genes, while limiting the unintended consequences frequently caused by gene knockout and overexpression.

## Article

The discovery of penicillin in the 1920s led to a new age of human and animal medicine as many antibiotics were quickly identified and developed, but the subsequent explosion of antibiotic treatments and applications has simultaneously driven microbial evolution and the development of widespread resistance^1,2^. Cell survival of antibiotic treatment due to antibiotic tolerance and persistence^3,4^ is a significant contributing factor to the abundance of antibiotic-resistant microorganisms. Persistence is a physiological state that enables cells to survive antibiotic treatment via temporary changes in phenotype, such as slowed growth and biosynthesis, rather than genotype (e.g. antibiotic resistance)^5^. Although persistence has been studied for over 70 years, there has been a lack of specificity in the literature between antibiotic tolerance and persistence^5,6^. Recently, a consensus statement that was released after a discussion panel with 121 researchers defined antibiotic persistence as a tolerant subpopulation of cells that result in a distinct phase of population decay^5^. We use population decay to differentiate between tolerance and persistence in this work (Fig. 1a).

**Fig 1.**
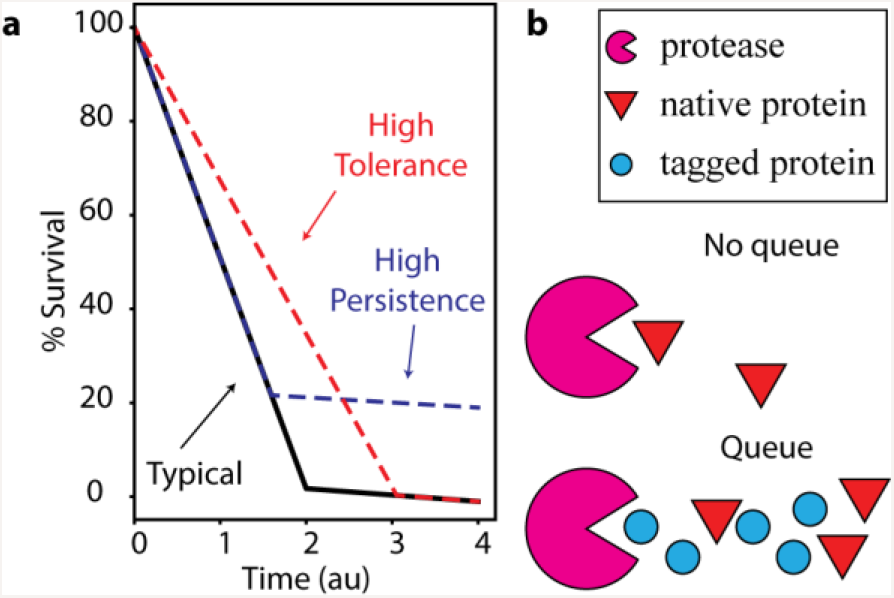
**a.** Examples of population decay in typical (black), high persistence (blue) and high tolerance (red) populations. A shift in tolerance can be distinguished from a change in the number of persisters. For example, a high persistence population can initially have the same decay rate as a typical population but have higher survival because of more persisters (dotted blue line). A high tolerance population can have the same persister level as a typical population but have a shift in the initial decay rate (dotted red line). **b.** A simple model of proteolytic queueing. When native proteins have low competition for the protease, there is no queue. Induction of synthetic tagged proteins competes with the native proteins for the protease and overloads the protease, which results in a proteolytic queue (bottleneck).

The widespread nature of persistence suggests that similar mechanisms exist to trigger the persistent state in prokaryotes. These mechanisms include many common systems, e.g. toxin-antitoxin (TA) systems and proteases. Although the precise role of TA systems in persistence is unclear, toxins in TA systems can trigger persistence when at a higher level than their cognate antitoxin^7–9^. Within the cell, the ratio of toxin to antitoxin is regulated during protein production^10–12^ and through degradation by proteases^13,14^. Proteases, such as Lon and ClpP, are largely responsible for protein degradation and cell maintenance^15,16^. They provide an essential level of protein regulation throughout the cell, including degradation of RpoS (a transcription factor that responds to stress)^17^ and tagged polypeptides (incomplete proteins) synthesized by stalled ribosomes that have been rescued by the trans-translation system^18^. In *E. coli*, *ssrA* (tmRNA) and *smpB* are the primary genes responsible for trans-translation, a cellular mechanism for recovering stalled ribosomes. A tmRNA molecule acts as a tRNA by binding to the A-site of a stalled ribosome. The ribosome is then transferred to translate the protein-coding region of the tmRNA, which adds an amino acid tag to target the polypeptide for degradation by ClpXP^18^. While *ssrA* is not essential in *E. coli*, *ssrA* knockouts cause growth defects, increase susceptibility to certain antibiotics^19^, and affect persistence^20,21^. Proteases and related chaperones are also consistently identified as persister related genes in gene knockout experiments^22,23^ and transcriptome analysis^24^. Indeed, a drug that targets persisters, acyldepsipeptide (ADEP4), activates the protease ClpP and lowers persister levels^25^. While most published articles focus on methods that reduce persister levels, conditions that increase their levels are integral to understanding the causative mechanisms of action and developing new drugs. As many persister studies incidentally examine antibiotic tolerance^5,6^, it follows that some of the above mechanisms may play a role in antibiotic tolerance.

Synthetic biology takes advantage of these mechanisms to develop new cellular circuits. For example, synthetic oscillators require rapid degradation of proteins, which is accomplished using the *ssrA* degradation tag^26–28^; the ssrA degradation tag is the amino acid sequence, AANDENYALAA^18^, which we abbreviate to LAA throughout. Previous work establishes that multiple circuits can be coordinated by overproduction of a common degradation tag to target proteins to a protease^29,30^. When a protease is overloaded, protein species compete for degradation; the enzyme is unable to keep up with the influx of new proteins^31^. This phenomenon can be explained by queueing theory, in which one type of customer competes for processing by servers, which has traditionally been applied to systems such as computer networks and call centers. Limited processing resources in a cell (e.g. proteases) cause biological queues^28,32^ (Fig. 1b). The queueing effect at the protease ClpXP is essential in allowing for oscillation of the highly used synthetic oscillator (often called Stricker oscillator or dual-feedback oscillator)^27,33^. Variations of this oscillator have been used in different strains of *E. coli*^27,29,30,34^, and in *Salmonella typhimurium*^35^, demonstrating that the queueing at ClpXP is not specific to one strain or species. The coupling of otherwise independent synthetic systems via proteolytic queueing demonstrates that queueing affects protein degradation and thus provides a tunable method of studying proteolytic degradation with little effect on cell growth^28–30,32^ compared to gene knockouts and overexpression of proteases^15,36,37^.

We set out to test the hypothesis that proteolytic queueing at the ClpXP complex effects survival of *E. coli* during antibiotic treatment. Previous studies have used knockout mutants to affect protease activity in *E. coli*, but these studies yielded mixed results^21,23,38,39^. The variability between results of knockout mutations could be due to differences in growth rates, which would modulate antibiotic efficacy. Proteolytic queueing is preferred over protease knockouts when probing antibiotic efficacy because protease knockouts often result in growth defects^15,36^, but proteolytic queueing does not noticeably affect cell growth or death^28–30,32^, even in stationary phase (Fig. S1). Our results show that during antibiotic treatment, degradation plays a role in cell survival and the effect is tunable using queue formation. Proteolytic queueing at ClpXP increases antibiotic survival and analysis of population decay with and without a queue demonstrates that queueing specifically increases antibiotic tolerance. We hypothesize that the queue is affecting the degradation of one or many regulatory molecules within the cell that cause downstream effects and enhance antibiotic tolerance. These results demonstrate that proteolytic queueing provides a new method to probe antibiotic tolerance and persistence.

## Results

### Proteolytic queueing affects tolerance

Cultures were grown to stationary phase and incubated for 24 hours prior to dilution into fresh media containing ampicillin to quantify persistence (see Methods). A proteolytic queue was induced via the production of a ssrA tagged fluorescent protein, CFP-LAA, expressed under an IPTG inducible promoter, Pl_ac/ara_-1. No apparent change in growth was observed by induction (Fig. S1) as reported previously^29,30^. The effects of queue formation on antibiotic survival are shown as the percentage of the population that survived ampicillin treatment (Fig 2). When CFP alone (the no degradation tag control) was overexpressed during ampicillin treatment, there was no significant effect on persister levels (p > 0.2, Fig. 2a). Queue formation (overexpression of CFP-LAA) during ampicillin treatment led to a 25-fold increase in survival after three hours in a concentration-dependent manner (Fig. 2b; p<0.0001, n ≥12).

**Fig. 2.**
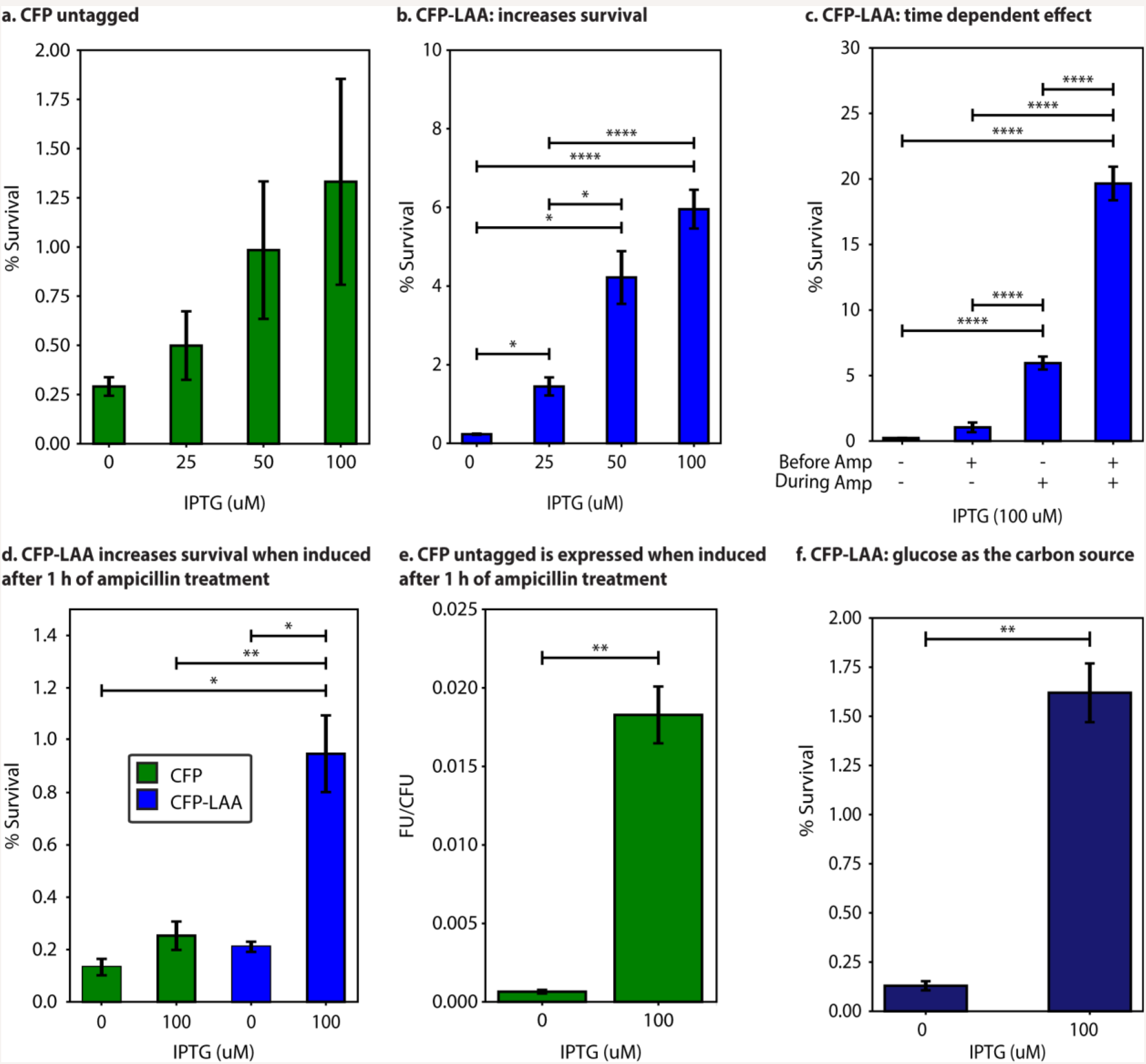
Proteolytic queueing affects survival of cells treated with the antibiotic ampicillin. **a.** Induction of untagged CFP during antibiotic treatment has no significant effect on survival (p>0.2). **b.** Induction of CFP-LAA during antibiotic treatment causes an increase in persistence. **c.** CFP-LAA was induced (+) with 100 μM of IPTG or not induced (−). Induction before ampicillin lasted 24 h in stationary phase prior to antibiotic treatment. Queueing only affects survival if the queue is maintained during ampicillin treatment. **d-e.** Expression of CFP or CFP-LAA was induced with IPTG one hour into the three-hour antibiotic treatment. Induction of CFP alone (no queue) had no significant effects on survival. Induction of CFP-LAA increased survival (**d**). Population fluorescence was measured for untagged CFP after antibiotic treatment, demonstrating that CFP is being produced via induction (**e**). **f.** Induction of CFP-LAA during antibiotic treatment causes an increase in persistence with glucose as a carbon source rather than glycerol, demonstrating that it is not a solely a carbon-specific phenomenon. Error bars represent SEM. n ≥ 3. *p<0.05. **p<0.01. ***p<0.001. ****p<0.0001.

When a queue was induced for 24 hours prior to ampicillin treatment the surviving population at three hours was over 80-fold higher than the uninduced population, only if induction was maintained during ampicillin treatment. However, if the inducer was removed during ampicillin treatment, the initial 24 hours of queueing had a minimal effect on survival at three hours (p>0.01, Fig. 2c). These results indicate that survival was affected by queue formation rather than CFP itself, and that the size of the queue (level and length of induction) determines the size of the effect. To confirm that these results are due to induction during antibiotic treatment, we waited one hour into ampicillin treatment before inducing expression of the fluorescent protein. As we previously observed, induction of untagged CFP had no apparent effect on persister levels (Fig. 2d), while quantification of fluorescence after ampicillin treatment confirmed that CFP was produced (Fig. 2e). Overexpression of CFP-LAA for two hours of ampicillin treatment still increased cell survival compared to the uninduced and untagged CFP populations (Fig. 2d).

We did further testing to confirm this effect is not specific to glycerol as a carbon source or ampicillin as the antibiotic. When glucose was the carbon source rather than glycerol, survival still increased due to CFP-LAA induction (Fig. 2f), which demonstrates that the effect is not directly related to the carbon source. We then tested the effects of queueing against the antibiotic ciprofloxacin, because ciprofloxacin targets DNA gyrase^40^ while ampicillin targets the cell wall^41^. CFP alone caused a slight increase in survival (Fig. 3a), however the CFP-LAA tag led to a 60- fold increase in survival (Fig. 3b).

**Fig. 3.**
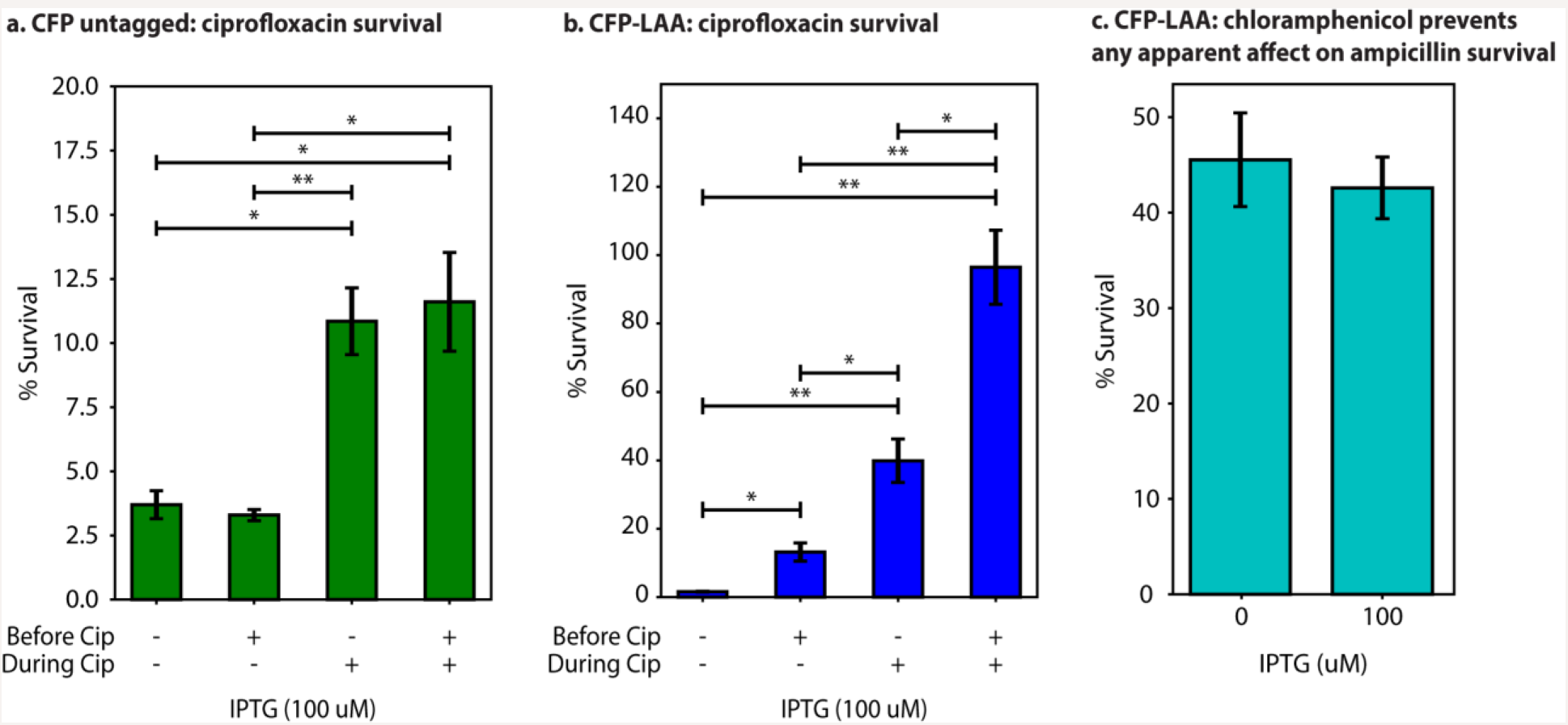
Proteolytic queueing effects in the presence of ciprofloxacin and chloramphenicol. **a.** Induction of untagged CFP during ciprofloxacin treatment increases survival less than 4-fold. **b.** Induction of CFP-LAA during ciprofloxacin treatment increases survival over 50-fold. **c.** Induction of CFP-LAA during ampicillin and chloramphenicol treatment has no apparent effect on persistence (p>0.7). X-axis labels correspond to Fig. 2. Error bars represent SEM. n≥3. *p<0.05. **p<0.01.

### Chloramphenicol inhibits the synthetic queue

Neither ampicillin nor ciprofloxacin directly affect production of the fluorescent protein (i.e. target transcription or translation) and thus should not prevent queue formation. On the other hand, an antibiotic that affects protein production should prevent queue formation, and therefore CFP-LAA induction would not affect survival in the presence of such an antibiotic. We found this to be the case when testing the effects of queueing on the survival of cells treated with chloramphenicol. Chloramphenicol is an antibiotic that inhibits protein translation by binding to bacterial ribosomes and inhibiting protein synthesis, thereby inhibiting bacterial growth^42^. Induction of CFP-LAA does not increase survival of antibiotic treatment when treated with chloramphenicol alone (Fig. S2), but chloramphenicol is not bactericidal, so we co-treated cultures with both ampicillin and chloramphenicol. The overall percent survival with chloramphenicol is much higher than with ampicillin alone, which is consistent with the literature^43^. As expected, co-treatment with ampicillin and chloramphenicol had no apparent effect on cell survival, supporting that even when CFP-LAA was induced the queue could not form if translation was blocked (Fig. 3c).

### Proteolytic queueing affects population decay

To gain further insight into the relationship between proteolytic queueing, tolerance and persistence, we measured how a proteolytic queue affects population decay by measuring survival for up to 8 hours of ampicillin treatment. Our results show a typical biphasic curve indicative of persister cells in the uninduced population. When the population is induced 24 hours prior to and during antibiotic treatment this curve shifts as the rate of population decay slows compared to uninduced cultures. The addition of the inducer exclusively during antibiotic treatment takes a similar effect between two and three hours into treatment. If the queue is induced 24 hours prior to antibiotic treatment, but the queue is not maintained (i.e. the inducer is removed during antibiotic treatment) the effect of the queue dissipates between one to two hours. There is no apparent difference between induced and uninduced cultures after 8 hours, which suggests there is little to no effect on persistence (Fig. 4a).

**Fig. 4.**
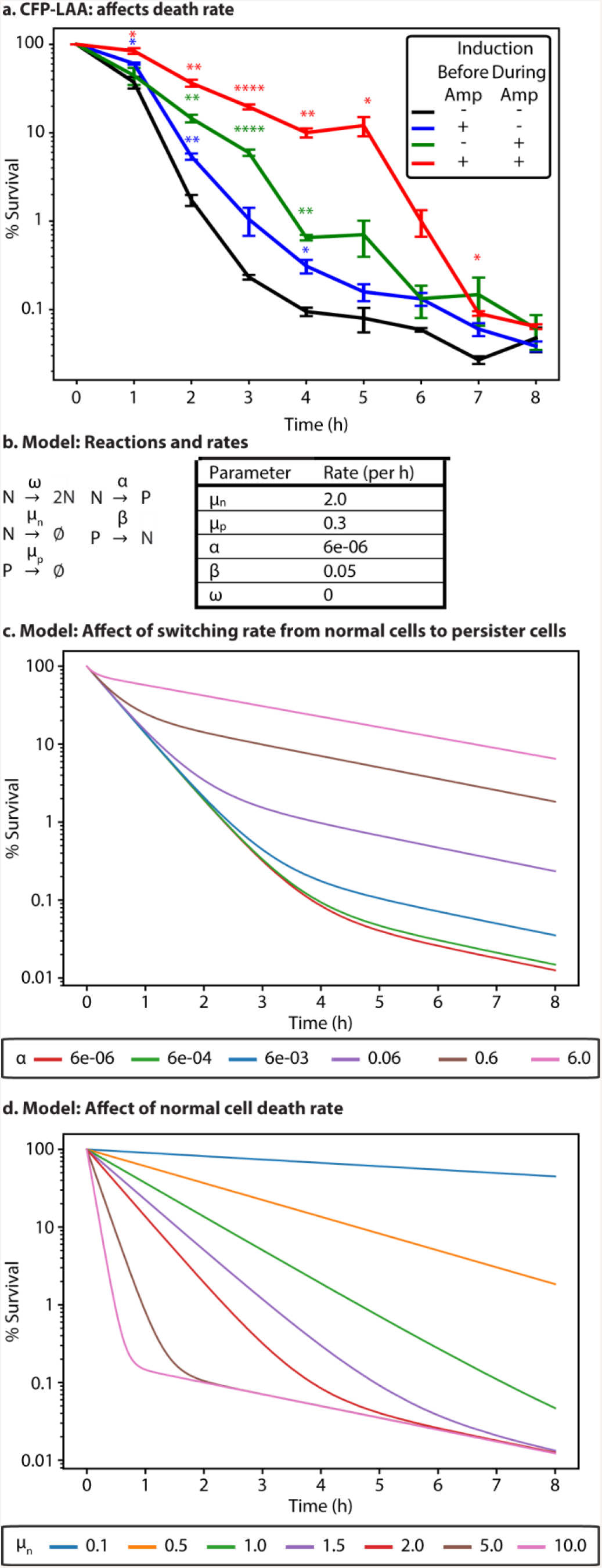
Time of queue formation influences survival. **a.** Stationary phase cells were diluted 1/100 into fresh media containing ampicillin (100 μg/ml) and sampled every hour for 8 h (n ≥ 3). Symbols (−/+) correspond to Fig. 2c. Error bars represent SEM. Asterisks indicate p-value (compared to no induction (black)) *p<0.05, **p<0.01, ***p<0.001, ****p<0.0001. There is 100% survival at time zero, because percent survival is determined based on the surviving CFU/ml compared to the CFU/ml at time zero. **b-d.** Stochastic model of population decay with antibiotic treatment. **b.** Reactions for the model (left) and baseline rates used for the simulations (right) unless stated otherwise (red lines below). Normal cell division (ω) was set to zero as dividing cells die during ampicillin treatment. **c.** Increasing the rate of entering persistence (α) increases cell number during the second phase of population decay. **d.** Decreasing the rate of normal cell death (μ_n_) causes the first phase of population decay to lengthen. **a,c,d.** Y-axes are in logarithmic scale.

### Computational modeling supports queueing-tolerance over queueing-persistence

Based on the *in vivo* results, we considered a computational model of population decay during antibiotic treatment modified from Kussel *et al.*^44^. In our model, the persister population (P) has a lower death rate than the susceptible population (N), where the death rates are represented by μ_p_ and μ_n_ respectively. We estimated μ_p_ and μ_n_ based on the experimentally determined decay rate of the uninduced population before and after two hours, and set the initial persister population to 0.2% of the total population (Fig. 4b). Normal (susceptible) cells enter persistence at rate α, and persister cells return to the normal state at rate β. The rates α and β were set relative to μ_n_ based on the relationship between these values in Kussel *et al*^44^. Our base model closely resembles population decay as measured in experimental tests. We use the model to determine whether the increase in overall population survival due to queue formation can be attributed to an increased rate of entering persistence (α) or increased tolerance (i.e. decreased μ_n_). Exploration of these parameters using stochastic simulations shows that increasing the rate at which normal cells become persisters (α) shortens the first phase of population decay and increases the number of persisters (Fig. 4c). Decreasing the rate of normal cell death (μ_n_) lengthens the first phase of population decay but has little to no effect on the number of persisters (Fig. 4d).

## Discussion

Proteolytic queueing is an integral component of native systems that has great potential for applications outside of synthetic biology. Here we show that queueing provides a tunable method to interfere with protease degradation and affect antibiotic tolerance. Increasing antibiotic tolerance in response to queueing was independent of the carbon source (glycerol or glucose) and antibiotic class (β-lactam or fluoroquinolone). When we prevented queue formation using chloramphenicol, adding IPTG did not affect survival of ampicillin. While CFP production alone slightly increased survival for ciprofloxacin, we suspect that high production of CFP with no apparent method of removal (besides cell division; minimal degradation) causes cell stress and affects survival, especially since high levels of fluorescent proteins can cause oxidative stress^45,46^, which is known to increase persistence^47–49^. However, because CFP-LAA is removed via degradation (indicated by lower fluorescence than CFP-untagged), the effects seen via overexpression of CFP should be less prominent during CFP-LAA overexpression. The results we describe here would not have been identified in a *clpP* knockout, because *clpP* knockouts have growth defects^37^ and any increase in tolerance would have been difficult to differentiate from this effect.

In some cases, the change in survival at three hours might be interpreted as a change in persistence; however, the shift in decay rates (as described in Fig. 1a) clearly demonstrates that queueing increases antibiotic tolerance rather than persistence. Furthermore, the effects caused by adding or removing the inducer during antibiotic treatment suggest that the change in antibiotic tolerance is due to an active response to the queue, which must be maintained to affect survival. Although persistence does not appear to be affected by the proteolytic queue at ClpXP, further overloading ClpXP is possible. Alternatively, the synthetic queue may not form in persister cells due to slowed translation and transcription. Our model supports that antibiotic tolerance is being affected rather than persistence, as altering survival of the ‘normal’ population more closely resembles the effects of proteolytic queueing than altering the rate of switching into persistence.

Proteolytic queueing is likely affecting the proteome of the cell, either directly or indirectly. Pleiotropic effects on protein content and gene regulation could be limiting antibiotic efficacy. Queue formation likely increases the intracellular concentration of multiple protein species causing a regulatory cascade. When considering proteins both degraded by ClpXP and related to persistence, TA systems are unlikely to be the causative factor, because decreasing degradation should increase antitoxin levels and decrease survival rather than increase survival as we observe. Regulatory proteins are possible candidates for the causative factor in queueing effects on tolerance. Such proteins include RpoS and DksA (both degraded by ClpXP), which have been implicated in persistence^21,49,50^ and may be involved in tolerance. Increased concentrations of these regulatory proteins due to slowed degradation could be causing downstream effects that lead to increased tolerance. In a similar vein, computational modeling has shown that altering degradation of MarA (a regulatory protein related to antibiotic tolerance) leads to increased coordination of downstream genes^51^. These and other regulatory proteins could be the cause of queueing-tolerance, as well as a yet unidentified protein(s). While these results are specific to queueing at ClpXP, tags are available to test the effects of queueing at other proteases (e.g. Lon and ClpAP)^32^.

The increase in antibiotic tolerance due to queue formation at ClpXP may be specific to overexpression of the LAA-tag, especially when considering that the number of LAA tagged proteins naturally increases during stress. The number of proteins with LAA tags increase during heat shock^52^, and queue formation at the proteases is likely a consequence of the increasing cellular traffic. If the native LAA tag is removed from SsrA while maintaining the ribosome rescue function, the survival of ampicillin treatment decreases in *E. coli*^21^. As the LAA tag could be a measurement of environmental stress, cells may have evolved to increase tolerance in response to increased queueing via LAA. Since ribosome rescue and proteolytic queueing are common across species, stress signaling via proteolytic queueing could be a general mechanism to regulate survival related genes. Identifying the key regulatory proteins in bacterial tolerance then understanding how these proteins interact are of great interest because they provide potential targets for killing bacterial pathogens, and proteolytic queues are a new method to explore these regulatory elements.

## Materials and Methods

### Strains and Plasmids

All strains are derived from *E. coli* DH5αZ1, and contain plasmids with the synthetic circuits, p24KmNB82 (CFP-LAA) and p24KmNB83 (untagged CFP) as described in REF^32^. DH5αZ1 was derived from *E. coli* K12 (arguably the most studied bacteria strain^53^), it is used by many in synthetic biology and outside the field^54–58^, this strain has previously been used to study persistence/tolerance or mechanisms related to them (e.g. toxin-antitoxin systems)^59–61^, and our previous queueing experiments used these derivitives^32^.

The cultures were grown in modified MMA media^62^, which we will refer to as MMB. MMB media consists of the following: K_2_HPO_4_ (10.5 mg/ml), KH_2_PO_4_ (4.5 mg/ml), (NH_4_)_2_SO_4_ (2.0 mg/ml), C_6_H_5_Na_3_O_7_ (0.5 mg/ml) and NaCl (1.0 mg/ml). Additionally, MMB+ consists of MMB and the following: 2 mM MgSO_4_ x 7H2O, 100 μM CaCl_2_, thiamine (10 μg/ml), 0.5% glycerol and amino acids (40 μg/ml). Cultures grown on glucose as the carbon source included 0.5% glucose instead of glycerol. Strains containing the plasmid p24Km and derivatives were grown in MMB+ kanamycin (Km, 25 μg/ml) or on Miller’s Lysogeny broth (LB) agar plates + Km (25 μg/ml). All cultures were incubated at 37°C and broth cultures were shaken at 250 rpm.

### Quantification of persistence

Persisters were quantified by comparing colony-forming units per milliliter (CFU/ml) before antibiotic treatment to CFU/ml after antibiotic treatment. The procedure for quantifying persister levels is based on previous research^59,63,64^ (Fig. S3). Briefly, overnight cultures were diluted 1/100 into fresh media and grown until they reach between OD_600_ 0.2-0.3. A reduced volume of culture (20 ml) was aliquoted into a 125 ml flask, and grown for 16 hours to enter stationary phase. Once in stationary phase, cultures were divided into two flasks with 0.2% arabinose, one flask of each replicate was also treated with 100 nM IPTG to induce expression under P_lac/ara-1_. Arabinose was added to both induced and uninduced cultures to maintain consistency (Fig. S4). All flasks were incubated for 24 hours before taking samples for plating and antibiotic treatment; cells were diluted 1/100^59,63^ into glass tubes, treated with 10X the MIC of ampicillin (100 μg/ml; Fig. S5) or 100X MIC of ciprofloxacin (1 μg/ml) at 37°C and shaken at 250 rpm for select time periods, 3 hours unless otherwise stated. Ampicillin solutions were stored at −80°C and only thawed once to reduce variability^19,65^. When indicated, samples were treated with chloramphenicol (5 μg/ml); cultures treated with chloramphenicol alone were diluted 1/10. Samples for quantification of CFU/ml were kept on ice and diluted using cold MMB before plating on LB/Km (25 μg/ml) agar plates. Cultures treated with ciprofloxacin were centrifuged at 16,000 *x g* for 3 minutes then washed with cold MMB to dilute ciprofloxacin before taking samples for quantification. LB agar plates were incubated at 37°C for 40-48 hours, then scanned using a flatbed scanner^66,67^. Custom scripts were used to identify and count bacterial colonies^68^ then calculate CFU/ml and persister frequency. Colonies were tested periodically for resistance, and we found no resistance in >350 colonies tested.

### Quantification of CFP

Cells were grown and treated with ampicillin as described in quantification of persistence above. After antibiotic treatment, 300 μl of cell culture was added to individual wells in a 96-Well Optical-Bottom Plate with Polymer Base (ThermoFisher) for fluorescence measurement using FLUOstar Omega microplate reader. The excitation and emission (Ex/Em) used for CFP measurement was 440/480. Readings were measured after four minutes of shaking to decrease variability between wells. Background fluorescence (mean fluorescence of MMB media) was subtracted from the raw reads. Fluorescence values were normalized by CFUs as determined by quantification of persistence, which was carried out simultaneously. Mean and SEM for fluorescence was determined across four biological replicates and three technical replicates.

### Computational modeling

Our model is modified from Kussel *et al.*^44^ where P is the persister population and N is the susceptible population (Fig 4b). Initial species counts P and N were set to 99800 and 200 respectively for all simulations, which we based on the percent survival of uninduced cultures. The death rate of N (μ_n_) and P (μ_p_) and the rate of entering (α) and exiting (β) persistence were set as shown in Fig. 4b unless otherwise stated. The rate of susceptible cell division (ω) was set to zero, as normal cells cannot divide without lysis during ampicillin treatment^69^. All simulations were performed using a custom implementation of the Gillespie algorithm^70^ in Python leveraging optimizations made possible by the Cython library^71^. Libraries from the SciPy stack^72^ were used for analysis.

### Statistics

All data is presented as mean ± SD or SEM of at least 3 biological replicates as appropriate^73^. Statistical significance for populations with the same number of replicates (n) was determined using one-way f-test to determine variance (p<0.001 was considered to have significant variance) followed by a Student’s t-test (no variance) or a Welch’s t-test (significant variance). Populations with different n values were compared using a Welch’s t-test. All statistical tests were run in Python using libraries from SciPy on groups with at least three biological replicates.

## Data availability

The data that supports the findings of this study are available from the corresponding author upon reasonable request.

## Code availability

Code used for model simulations is available on GitHub at https://github.com/ctogle/mini_gillespiem. Code used for colony counting is available on GitHub at https://github.com/hdeter/CountColonies.

## Acknowledgments

This work is supported by the Hatch project grant no. SD00H653-18/project accession no. 1015687 from the USDA National Institute of Food and Agriculture.

## Author contributions

H.S.D wrote the manuscript, developed custom code for colony counting, and ran statistical analyses. A.A. performed ampicillin and ciprofloxacin persister assays. P.J. performed plate reader assays. E.S. performed chloramphenicol persister assay. C.T.O. and H.S.D. adapted the persister model and ran stochastic simulations. N.C.B. initiated and directed the project. All authors contributed to discussing and editing the manuscript.

## Competing interests

The authors declare no competing interests.

**Correspondence and request for materials** should be addressed to N.C.B.

## Supplementary Figures and Data

**Fig S1.**
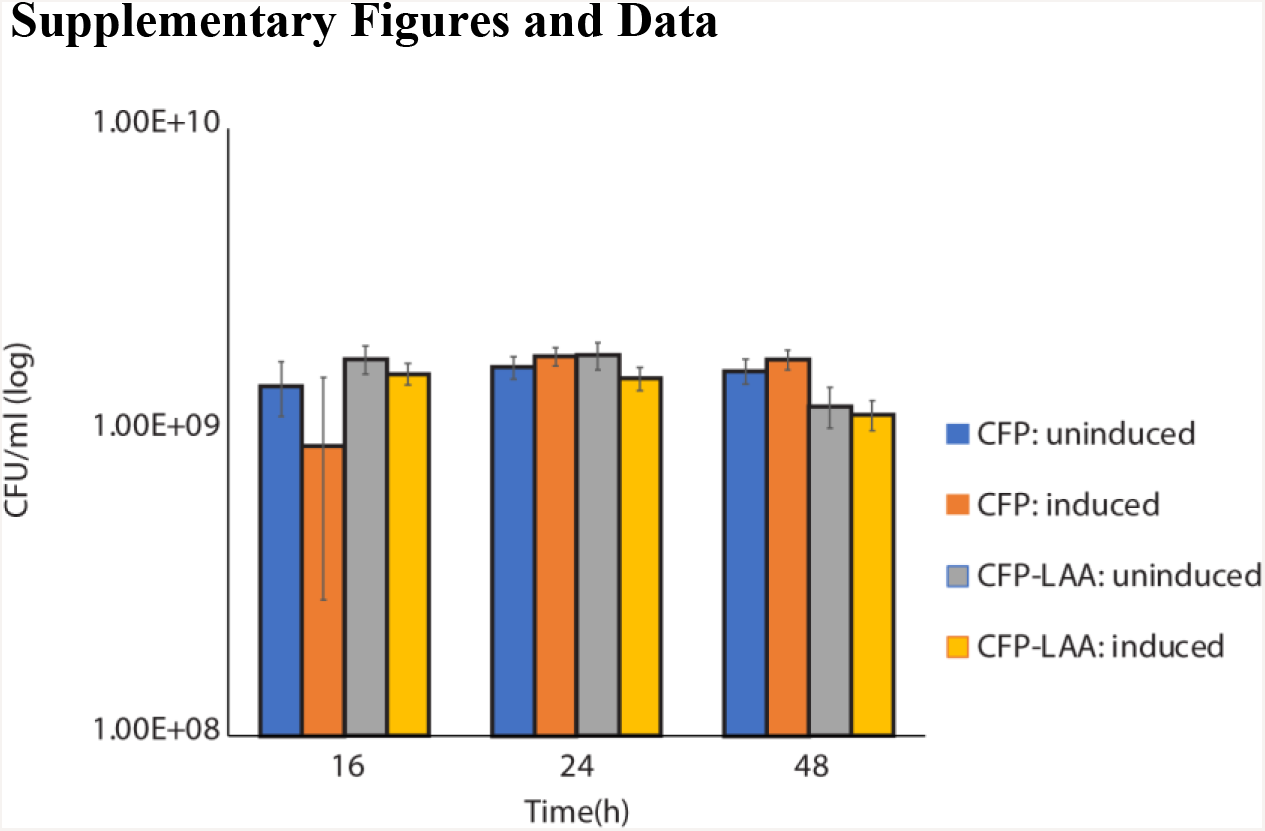
Induction of untagged CFP and CFP-LAA tag has no apparent effect on growth in MMB+ media. The Y-axis is in log CFU/ml of induced and uninduced cultures over 48 hours. n≥3. Error bars represent the standard deviation.

**Fig. S2.**
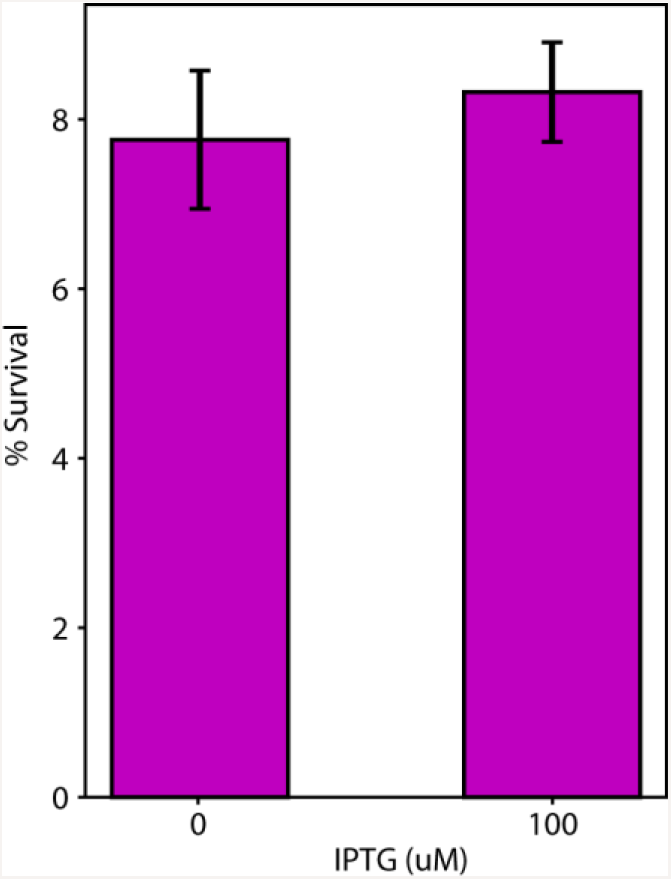
Induction of CFP-LAA does not increase survival of cells treated with chloramphenicol. Cultures were treated with chloramphenicol, an antibiotic that inhibits translation, after a 1/10 dilution into fresh media from stationary phase. Induction of CFP-LAA via IPTG had no significant change in persistence compared to the uninduced cultures (p>0.7; n≥3). Error bars represent SEM.

**Fig. S3.**
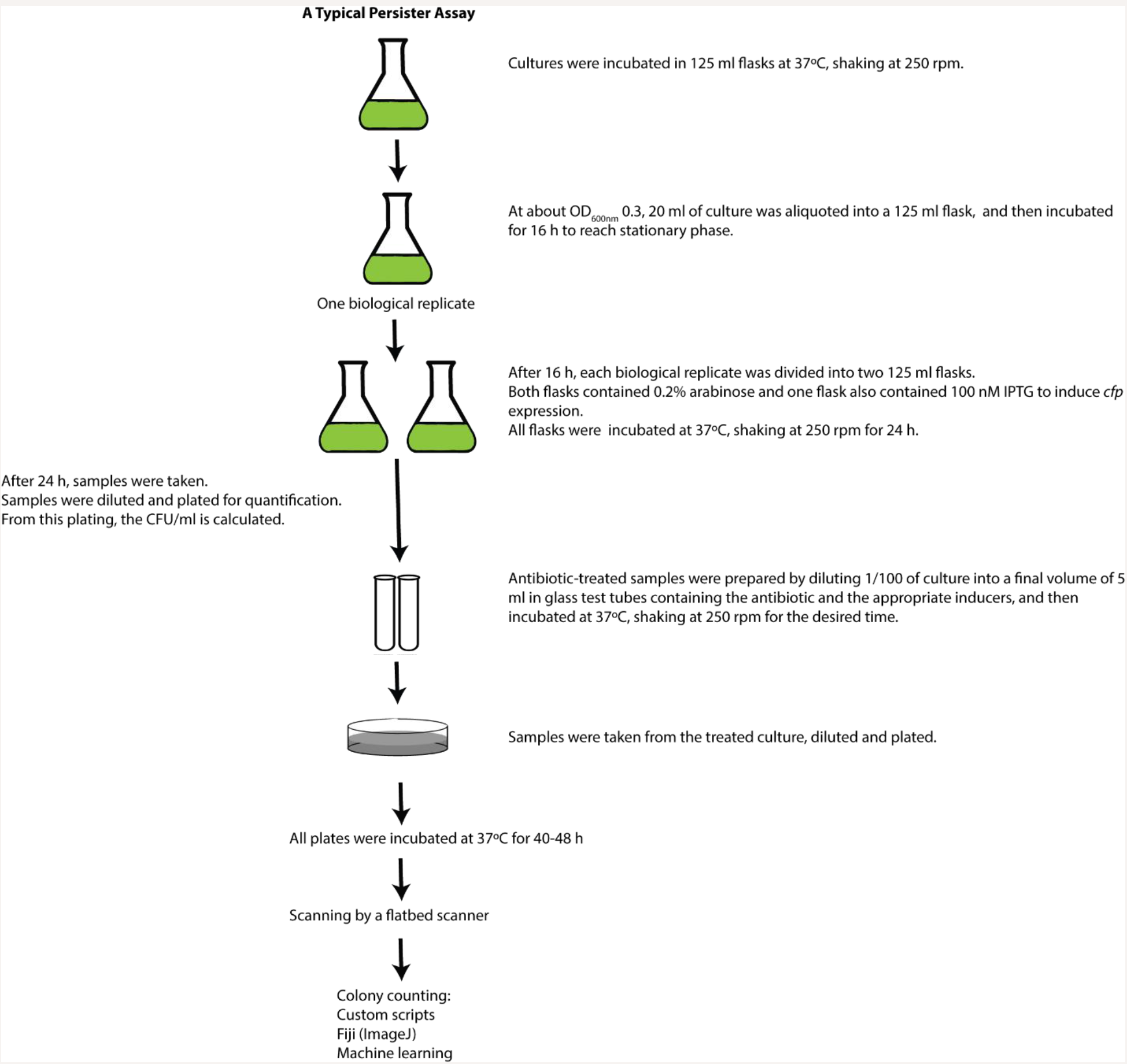
Persister assay flow chart. See Methods for details.

**Fig. S4.**
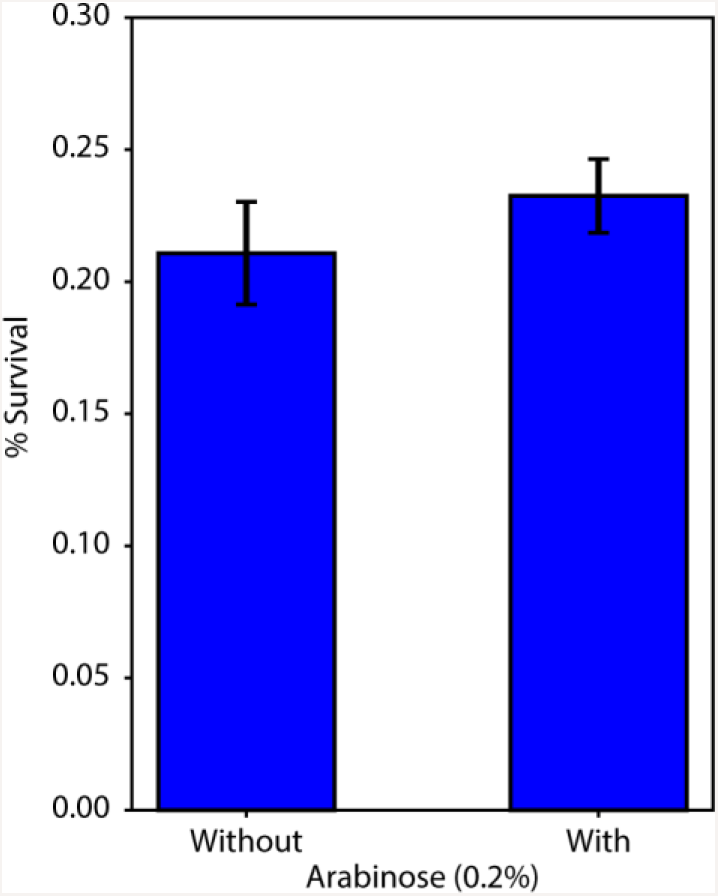
The addition of arabinose had no apparent effect on the tolerance/persister level during ampicillin treatment. Both IPTG and arabinose are inducers for CFP untagged and CFP-LAA tagged proteins. IPTG induces expression, arabinose alone does not induce expression, but arabinose can enhance expression when used in combination with IPTG. The effect of adding arabinose (0.2%) on tolerance/persistence to ampicillin was tested with CFP-LAA. Adding arabinose does not have a significant effect on survival of cells after 3 hours of ampicillin treatment (p>0.3). Error bars represent SEM. n ≥ 3.

**Fig S5.**
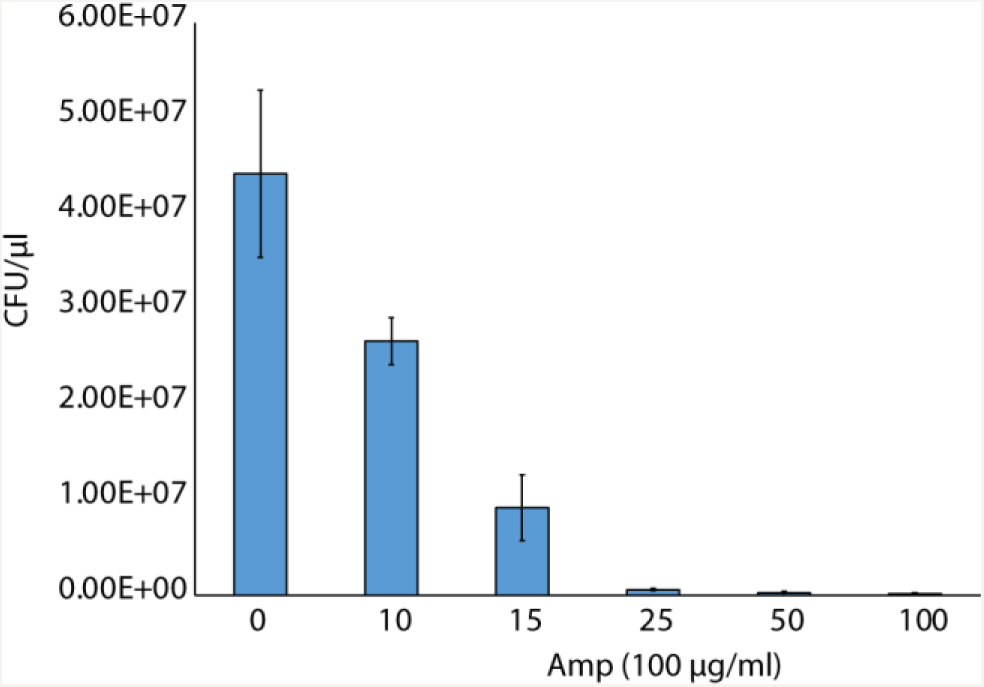
Determination of Minimal Inhibitory Concentration (MIC) for ampicillin. Exponential phase cultures were treated with different concentrations of ampicillin. The MIC was determined to be 10 μg/ml (p <0.03 compared to zero). Error bars represent the standard deviation.

